# Reversal of contractile defects by mediating calcium homeostasis in human mini-heart models of heart failure with preserved ejection fraction (HFpEF) leads to first-in-human gene therapy clinical trial

**DOI:** 10.1101/2024.08.27.609034

**Authors:** Kevin D. Costa, Andy O. T. Wong, Suet Yee Mak, Erin G. Roberts, Wendy Keung, Claudia Correia, Anna Walentinsson, Jonas Christoffersson, Alice Cheung, Deborah K. Lieu, Karin Jennbacken, Qing-Dong Wang, Roger J. Hajjar, Ronald A. Li

## Abstract

**Aims:** Heart failure with preserved ejection fraction (HFpEF), is a global health problem lacking disease-modifying therapeutic options, reflecting a lack of predictive models for preclinical drug testing. Aligned with FDA Modernization Act 2.0, we aimed to create the first *in vitro* human-specific mini-heart models of HFpEF, and to test the efficacy of a candidate gene therapy to improve cardiac kinetics and correct the disease phenotype.

**Methods and Results:** Healthy human pluripotent stem cell-derived ventricular cardiomyocytes were used to bioengineer beating cardiac tissue strips and pumping cardiac chambers. When conditioned with transforming growth factor-β1 and endothelin-1, these mini-heart models exhibited signature disease phenotypes of significantly elevated diastolic force and tissue stiffness, and slowed contraction and relaxation kinetics, with no significant deficit in systolic force or ejection fraction versus unconditioned controls. Bioinformatic analysis of bulk RNA sequencing data from HFpEF mini-heart models and patient ventricular samples identified downregulation of SERCA2a of the calcium signalling pathway as a key differentially expressed gene. After dosage optimization, AAV-mediated expression of SERCA2a abrogated the disease phenotype and improved the cardiac kinetics in HFpEF mini-Hearts.

**Conclusions:** These findings contributed to FDA approval of an ongoing first-in-human gene therapy clinical trial for HFpEF, with Fast Track designation. We conclude that such human-based disease-specific mini-heart platforms are relevant for target discovery and validation that can facilitate clinical translation of novel cardiac therapies.

**Translational Perspective:** Heart failure with preserved ejection fraction (HFpEF) is a significant and growing global health concern lacking disease-modifying therapeutic options, reflecting inadequate preclinical models of the disease. Aligned with FDA Modernization Act 2.0, we created the first *in vitro* human-specific mini-heart models of HFpEF, demonstrated phenotypic disease characteristics of elevated stiffness and slowed kinetics, showed transcriptomic consistency with HFpEF patient data, identified SERCA2a as a key downregulated gene, performed dosing titration of SERCA2a gene therapy, and showed improvement of cardiac kinetics post-treatment. The findings contributed to FDA approval of an ongoing first-in-human gene therapy clinical trial for HFpEF.

## Introduction

Heart failure with preserved ejection fraction (HFpEF) is becoming the most prevalent cause of heart failure,^1-3^ accounting for over 50% of all HF cases. Unlike other cardiovascular diseases, the prevalence of HFpEF is increasing in Europe and the US, with a substantial economic burden, and high mortality comparable to heart failure with reduced ejection fraction (HFrEF).^1-4^ Unfortunately, many therapies that reduce mortality and morbidity in HFrEF have little benefit for HFpEF patients because the biology and clinical course for the two diseases are very different. Despite a left ventricular (LV) ejection fraction of 50% or more, the main clinical presentation of patients with HFpEF is exercise intolerance and dyspnea even at very low levels of exertion,^5^ leading to a downward spiral of inactivity and deconditioning that compounds the underlying cardiac disease. The fundamental structural and functional characteristics of the HFpEF heart include delayed and incomplete LV relaxation and reduced LV compliance.^6, 7^ Two pathological characteristics markedly prevalent in patients with HFpEF are cardiac myocyte hypertrophy and cardiac fibrosis, as confirmed in a large, prospective analysis of HFpEF myocardial tissue.^8, 9^

There is an urgent need to develop targeted therapies for this largely untreated patient population. Although several experimental animal models have been established and reported to mimic aspects of HFpEF,^10-13^ they fail to recapitulate key characteristic clinical manifestations due to species-specific and other inherent differences.^14, 15^ Various lines of evidence from these models hint at the sarcoplasmic reticulum (SR) Ca^2+^-ATPase pump (SERCA2a), crucial for calcium homeostasis, as a therapeutic target for HFpEF, but direct human relevance has not been demonstrated.

Bioengineered cardiac tissues, created from human pluripotent stem cell (hPSC)-derived cardiomyocytes, have successfully modeled aspects of native human heart structure and function,^16^ including cardiotoxic and therapeutic responses,^17, 18^ and phenotypic characteristics of genetic and acquired cardiomyopathies.^19, 20^ Indeed, the recently passed FDA Modernization Act 2.0 ends the long-standing animal mandate and promotes the use of validated *in vitro* human alternatives for next-generation drug development.^21^Here, our objective was to create the first *in vitro* human heart models of HFpEF, and to apply them for therapeutic target discovery and validation. We report the use of our established mini-heart platform,^17^ which was developed in recognition of the need for human-specific cardiac assays to improve the success of clinical trials. These include the trabecular muscle-like human ventricular cardiac tissue strip (hvCTS) assay for measuring cardiac contractility,^16, 17^ and the miniature heart-like human ventricular cardiac organoid chamber (hvCOC) assay for measuring clinically relevant metrics of ventricular pump function.^22^ Based on the known biology of HFpEF, hvCTS and hvCOC mini-hearts were treated to induce key disease characteristics of HFpEF, followed by testing the functional consequences of a gene therapy approach. These data have contributed to our successful Investigational New Drug (IND) and Fast Track Designation (FTD) applications to the U.S. FDA, with an ongoing clinical trial.

## Methods

### Induction of HFpEF in human ventricular cardiac tissue strips (hvCTS)

Three-dimensional (3D) multicellular human ventricular cardiac tissue strips (hvCTS) were engineered as previously described.^16, 23^ Briefly, human pluripotent stem cells from either of two established healthy female cell lines that were obtained commercially for research purposes (HES2: NIH code ES02, from ES Cell International; or H7: NIH code WA07, from WiCell) were directed to differentiate into beating cardiomyocytes with high ventricular-specificity (hvCMs), using a published small molecule-based embryoid body method.^24^ The percentage of cells expressing cardiac troponin T was verified to be between 65% and 85% using flow cytometry on differentiation day 15, with the remainder being cardiac fibroblast-like stromal cells. Cardio-clusters that satisfied this verification criteria were then dissociated into small cell clusters and allowed to recover in the incubator for 3 days before hvCTS construction. Each hvCTS consisted of 1.3 × 10^6^ hvCMs and 1.3 × 10^5^ human foreskin fibroblasts in a 100-μL ice-cold solution of 2 mg/ml collagen I, 0.80–0.95 mg/mL Matrigel, 0.6X PBS, 20 mM NaOH, 0.8X Minimum Essential Medium (MEM, Sigma-Aldrich), 1.6 mM HEPES, and 0.1X hvCTS maintenance medium (see composition below). A volume of 100 μl of the final cell-collagen mixture was then added to each polydimethylsiloxane (PDMS) bioreactor, consisting of a rectangular well with a force-sensing cantilever post at each end, and returned to the incubator to form the trabecular muscle-like hvCTS attached between the two end-posts. The hvCTS were maintained in Dulbecco’s Modified Eagle Medium (DMEM) medium supplemented with 10% newborn calf serum (NCS; Gibco), with daily half-medium changes for 5 days to allow tissue formation and compaction. To induce the HFpEF phenotype, randomly selected hvCTSs were treated with TGF-β1 (1 ng/mL) for 4.5 days, followed by treatment with TGF-β1 (1 ng/mL) plus ET-1 (100 nM) for 1 additional day prior to contractile assessment. Time-matched hvCTS without TGF-β1/ET-1 treatment served as healthy controls.

Dose response effects for separate treatments of TGF-β1 (0-30 ng/mL) or ET-1 (0-100 nM) were also tested to identify the concentrations that could elicit a significant change in stiffness without compromising the integrity and success rate of the treated tissues, yielding optimized concentrations (1 ng/mL TGF-β1 and 100 nM ET-1) for effective combined induction of HFpEF models.

### Measurement of force, stress, strain, and stiffness in hvCTS

Force generated by the hvCTS was measured at 37°C in phenol red-free DMEM medium with HEPES buffer (1.8 mM Ca^2+^) using an automated force measurement system (CTScreen, Novoheart) that records displacement of the PDMS cantilever end-posts, captured in real time with a high-speed (100 frames per second) camera (Allied Vision, Exton, PA) and custom LabVIEW software (National Instruments, Austin, TX) under spontaneous beating (unpaced) or electrically paced conditions (field stimulation at 1.0, 1.5, 2.0, 2.5 and 3.0 Hz). A beam-bending equation was then used to calculate twitch force, as described previously.^25^ For isometric force assessment, the hvCTS was removed from the PDMS bioreactor and attached to the force transducer and stepper motor of an isometric muscle bath system (Aurora Scientific). Diastolic, systolic, and developed force and other contractile parameters were acquired at 37°C under spontaneously beating and electrically paced conditions (as above) at 0-50% strain at 5% intervals. Stress was calculated as force normalized by the tissue cross-sectional area, and passive stiffness (i.e., secant modulus) was calculated as diastolic stress divided by strain at a given percent strain. Data analysis was performed by research assistants blinded to the experimental group assignments.

### HFpEF induction in human cardiac organoid chambers (hvCOC)

Three-dimensional, pumping human ventricular cardiac organoid chamber (hvCOC) mini-hearts were engineered as previously described,^22^ with an ultra-compliant indwelling elastomer balloon.^26^ Briefly, each hvCOC consisted of 1.0 × 10^7^ hvCMs and 1.0 × 10^6^ human foreskin fibroblasts in a 1650-μL ice-cold solution of 2 mg/mL collagen I, 0.80–0.95 mg/mL Matrigel, 0.6X PBS, 20 mM NaOH, 0.8X MEM, 1.6 mM HEPES, and 0.1X hvCTS maintenance medium (defined above). The cell-collagen mix was added to the space between a cup-shaped agarose mold and a concentric elastomer balloon, with a porous polyethylene ring placed just above the base of the balloon but submerged in the cell suspension to enhance tissue attachment. The bioreactor was incubated for 1 hour to allow gelation of tissue before filling with NCS medium (8 mL). Medium was changed every 24 hours while the hvCOC was compacting, and every other day for 5 days after the hvCOC was removed from the agarose mold. To induce the HFpEF phenotype, randomly selected hvCOCs were treated with the same protocol of TGF-β1 (1 ng/mL) for 4.5 days and then TGF-β1 (1 ng/mL) plus ET-1(100nM) for an additional day. Time-matched hvCOC without TGF-β1/ET-1 treatment served as healthy controls.

### Measurement of stiffness and function in hvCOC

As detailed elsewhere,^22^ pump function was evaluated using a commercial automated system (COScreen, Novoheart) that includes a high-sensitivity pressure catheter (Millar) advanced into the lumen of the hvCOC chamber for pressure measurements. A digital camera (Prosilica GX1050C, Allied-Vision) was mounted outside of the hvCOC bioreactor and permitted direct tissue monitoring for determination of chamber area profile. Chamber pressure and digital video were acquired simultaneously under spontaneously beating and electrically paced conditions (as above) at 0, 25, 50, and 100 μL of volumetric loading. Stiffness was calculated from the slope of the change in diastolic pressure versus change in diastolic area of the chamber resulting from volumetric loading.

### Bulk RNA sequencing of HFpEF hvCTS and hvCOC

Frozen TGFb1/ET1-treated and untreated hvCTS (n=7 per condition) and hvCOC (n=5 per condition) were lysed using Agencourt RNAdvance Tissue kit (Beckman Coulter) lysis buffer and homogenized at 1000 rpm for 2 min with 5 mm Stainless Steel Beads (Qiagen) using 1600 MiniG Tissue Homogenizer (SPEX SamplePrep). Total RNA was extracted using Agencourt RNAdvance Tissue kit including DNase treatment for 25 min at 37°C (Ambion DNaseI, ThermoFisher) on an automated Biomek i7 Hybrid system (Beckman Coulter). RNA quantification and quality control was performed on a 48-capillary 5300 Fragment Analyzer System (Agilent). Total RNA-seq library preparation was performed using KAPA RNA HyperPrep Kit with RiboErase (Roche) with 450 ng of input RNA, 5 min fragmentation at 95°C, and 9 PCR cycles performed on a Tecan Fluent 1080 system (Tecan). Resulting libraries were quantified on a 48-capillary 5300 Fragment Analyzer System and sequenced with a 100 bp paired-end setting on a v1.5 S1 NovaSeq6000 kit (Illumina).

Read quality for all libraries was assessed using FastQC (v0.11.7), Qualimap (v2.2.2c) and samtools stats (v1.9). Quality control (QC) metrics for Qualimap were based on a STAR (v2.7.2b) alignment against the human genome (GRCh38, Ensembl v100). Next, QC metrics were summarized using MultiQC (v1.9). Sequencing adapters were then trimmed from the remaining libraries using NGmerge (v0.3). A human transcriptome index consisting of cDNA and ncRNA entries from Ensembl (v100) was generated and reads were mapped to the index using Salmon (v1.1.0). The bioinformatics workflow was organized using Nextflow workflow management system (v20.10) and Bioconda software management tool.

Differential expression gene (DEG) analysis between HFpEF-hvCOC/hvCTS models (TGF-β1/ET-1-treated versus control) was performed using DESeq2 (v1.24.0), using percentage of cardiac troponin T-expressing cells before tissue fabrication and age of tissue at time of data collection as baseline covariates. Read quality for all libraries was accessed using FastQC. Sequencing paired-end reads were mapped towards the human reference genome (GRCh38) using Star Aligner.

A combined criteria of FDR ≤ 0.05 and the log2 fold change |log2(FC)| ≥ 0.5 was used to define the DEG sets for downstream analysis. The DEGs were analysed through the use of IPA (Ingenuity Pathways Analysis; QIAGEN Inc.). This tool uses the information in the Ingenuity® Knowledge Base to assess signalling and metabolic pathways, upstream regulators, regulatory effect networks, and disease and biological functions that are likely to be perturbed based on a data set of interest. The IPA analysis was performed to identify significantly dysregulated canonical pathways, disease and biofunctions, and cardiotoxicity functions in HFpEF-hvCOC/hvCTS (TGF-β1/ET-1-treated) group versus control (non-treated) group. The enrichment was tested using Fisher’s exact test with p-value ≤ 0.05. Activation Z-score was used to predict activated or inhibited functions/pathways.

DEGs from the HFpEF-hvCOC/hvCTS models were compared to published human heart failure transcriptomic data^8^ (RNA sequencing data from right ventricular septal biopsies of male and female HFpEF patients undergoing invasive hemodynamic testing (n=41) vs. brain-dead organ donor control hearts (n=24)), both on gene-level and by comparative functional enrichment analyses using IPA.

### AAV1-SERCA2a transduction of hvCTS and hvCOC

SERCA2a mRNA was delivered by transduction with GMP-grade AAV1-SERCA2a viral vector (lot number S827002, from Sardocor, Medera Inc., Boston, MA) comprising an expressible coding region for SERCA2a. The AAV1-SERCA2a drug product was suspended in culture media and delivered to dissociated human pluripotent stem cell-derived hvCMs at day 15 post-differentiation at 1×10^2^ to 1×10^5^ viral genomes (vg) per cell. The hvCMs were then fabricated into hvCTS or hvCOC and treated with HFpEF induction medium containing TGF-β1/ET-1, as described above. The human tissue models were maintained in such medium until testing. Treatment control studies used AAV1-GFP (Virovek, Houston, TX) or non-transduced (NT) cells. SERCA2a gene expression level was evaluated by RT-qPCR analysis of isolated mRNA.

### Statistical analysis

Data are presented as mean ± SEM unless otherwise specified. Depending on the number of groups and factors being compared, statistical analysis was performed using Student’s two-tailed t-test; one-way ANOVA followed by Dunnett’s multiple comparison test; or two-way ANOVA followed by Sidak’s multiple comparison test. Statistical significance was accepted for values of p ≤ 0.05.

## Results

During the last decade, significant effort has focused on the elucidation of signaling pathways that mediate the complex response of cardiomyocytes to hypertrophic stimuli and the progression to HF.^27-35^ Endothelin-1 (ET-1) is a potent stimulator of cardiac myocyte hypertrophy, and is elevated in patients with HFpEF.^36^ Cardiac fibrosis (CF) is also highly associated with heart failure (HF) including HFpEF, where it has been proposed to play a key role in the progression of the disease,^37^ and CF has been associated with mortality in patients with HFpEF.^38^ In particular, members of the Transforming Growth Factor-β family (e.g., TGF-β1), which are secreted in the cardiac interstitium, influence the activation of specific aspects of the fibrotic response,^9, 10, 39-41^ impacting the synthesis, processing and metabolism of the extracellular matrix. This foundation of knowledge informed our strategy to model the cardiac phenotype of HFpEF *in vitro*.

### hvCTS tissue strip model of HFpEF exhibits elevated stiffness and impaired contractile kinetics

The hvCTS models were fabricated as previously detailed^16, 17^ using healthy hPSC-derived ventricular cardiomyocytes (hvCMs) and fibroblasts suspended in a collagen-based hydrogel to form a thin strip of cardiac muscle in a custom bioreactor with integrated end-posts for automated measurements of twitch force and contractility. Given the well-defined biological roles of TGF-β1 and ET-1 in HFpEF,^36-38^ we tested the combination of the pro-fibrotic signal and hypertrophic stimulator to model the HFpEF phenotype *in vitro* by examining the functional consequences of their combined effects on the contractility and stiffness of hvCTS. **Figure 1A** shows representative images of time-matched hvCTS under Control conditions and after combined TGF-β1/ET-1 treatment over 5.5 days. The end-posts for the treated group were strongly deflected even in the relaxed (diastolic) state, indicating higher passive tension of the attached hvCTS (**Fig. 1C**). Consistently, the TGF-β1/ET-1-treated group displayed significantly slower contraction and relaxation kinetics compared to Control (**Fig. 1B,C**; n = 4-20, **p<0.01 and ***p<0.001). The hvCTS were also tested using a physiologic muscle bath system with isometric length-control during uniaxial stretch. Measurements of passive stiffness, developed force, maximum +dF/dt and maximum -dF/dt versus percent strain (**Fig. 1D**) collectively showed that TGF-β1/ET-1-treated hvCTS displayed significantly increased passive stiffness along with slowed contraction and relaxation kinetics compared to Controls, with no significant reduction in systolic or developed force, which are key characteristics observed in the left ventricles of HFpEF patients. We note that single-factor treatments of hvCTS with either TGF-β1 or ET-1 individually did not elicit the characteristic combination of elevated stiffness and depressed kinetics achieved with the combination treatment (**Supplemental Figs. 1 and 2**).

**Figure 1.**
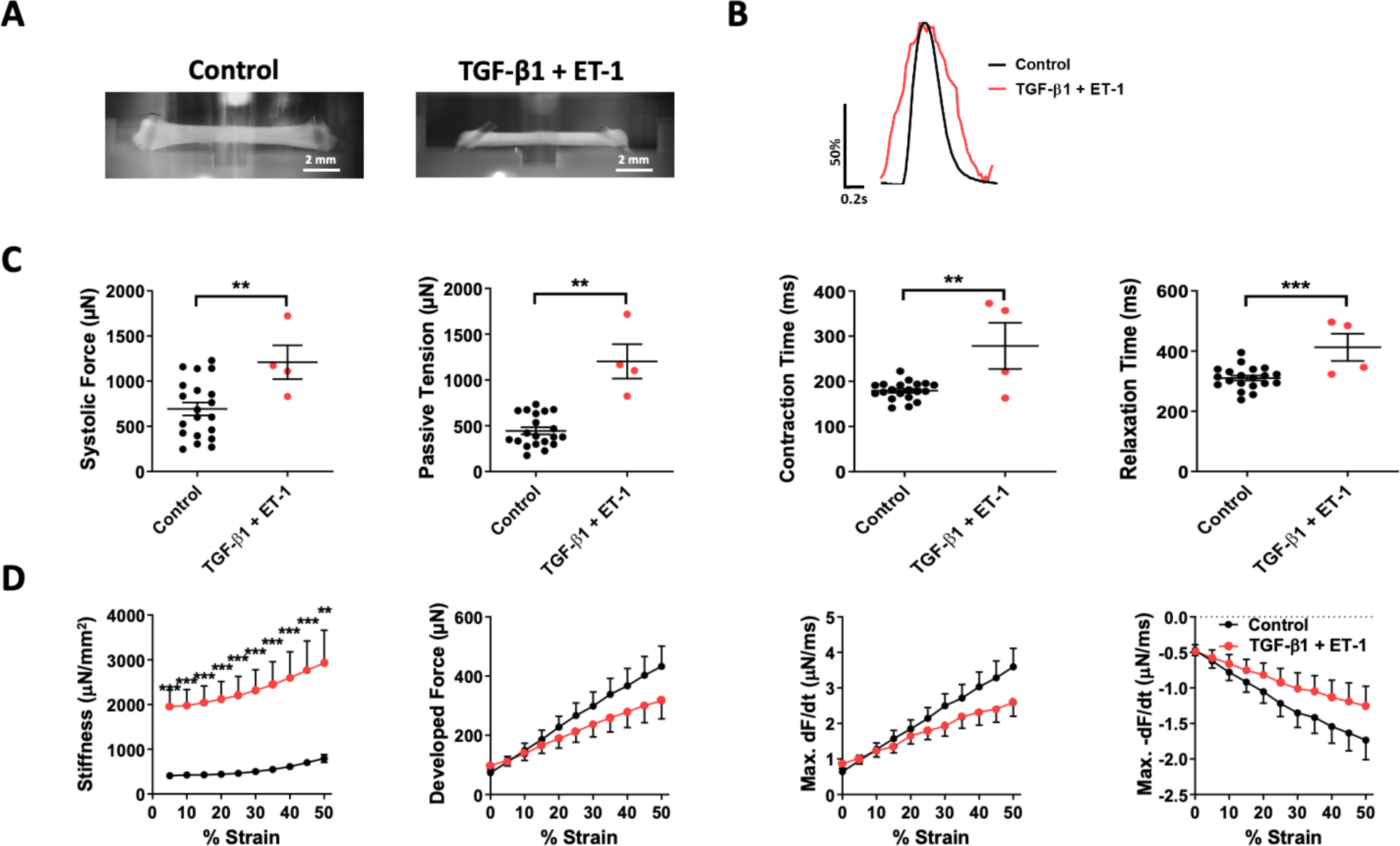
Functional consequences of combined effects of TGF-β1/ET-1 on contractility of human ventricular cardiac tissue strips (hvCTS). (A) Representative images of time-matched (Day 10) hvCTS at a relaxed diastolic state under control conditions and with combined Transforming Growth Factor-β1/Endothelin-1 (TGF-β1/ET-1) treatment. (B) Representative normalized single-twitch force tracings at 1.0-Hz pacing show slower contraction and relaxation kinetics in the TGF-β1/ET-1-treated group (red) than the control group (black). (C) Dot plots for systolic force, passive tensile force, contraction time, and relaxation time recorded by post-tracking measurements during 1.0-Hz field stimulation, showing mean ± SEM for control group (black) and TGF-β1/ET-1-treated group (red). n = 4-20 tissues; Student’s t-test. (D) Stiffness, developed force, max +dF/dt and max -dF/dt versus percent strain for control (black) and TGF-β1/ET-1-treated (red) hvCTS, recorded using a physiologic muscle bath system with isometric length-control. n=7-12 tissues; mean ± SEM; Two-Way ANOVA followed by Sidak’s multiple comparison test. **p<0.01 and ***p<0.001.

**Figure 2.**
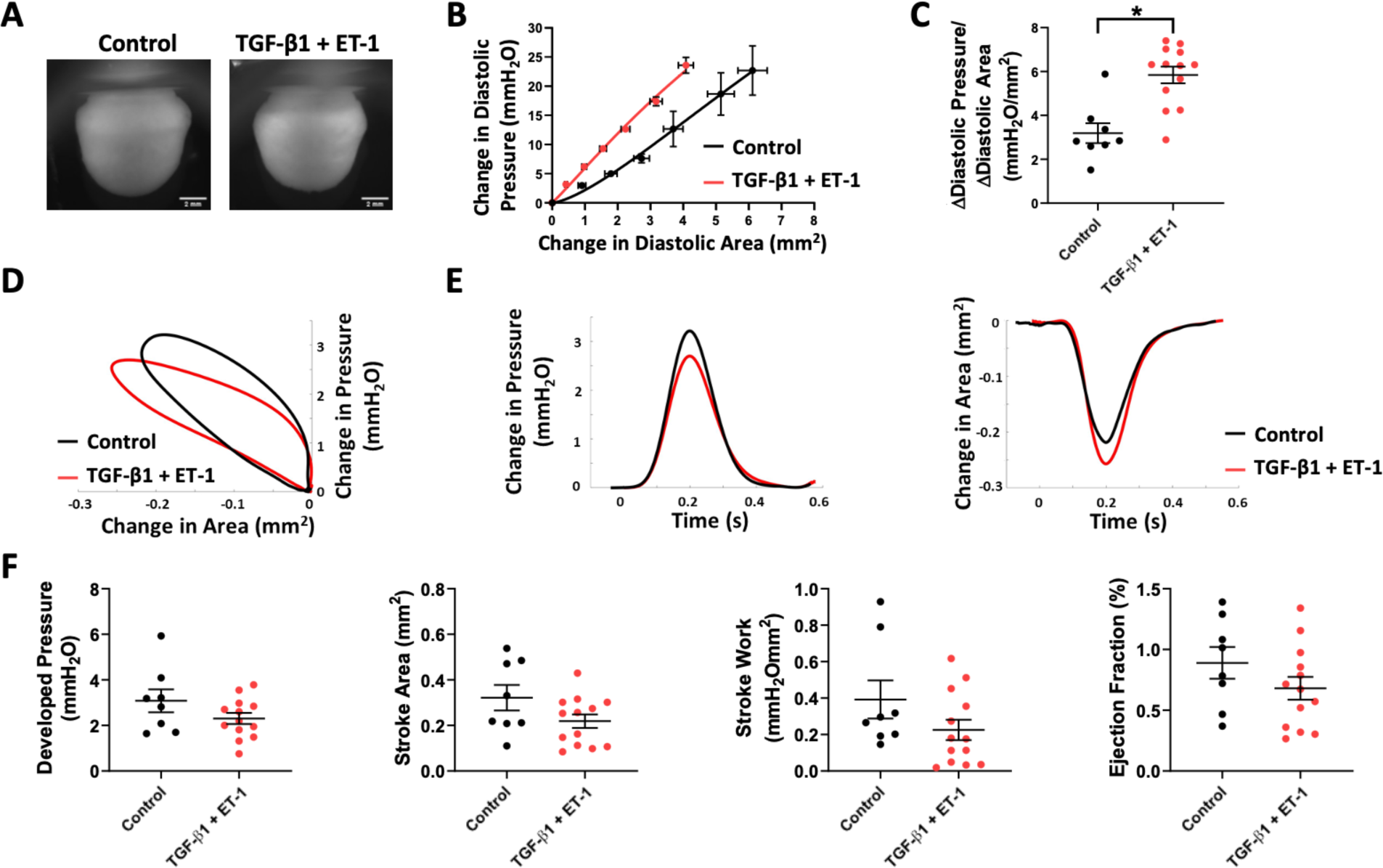
Combined TGF-β1/ET-1 treatment conferred HFpEF phenotypic characteristics on human ventricular cardiac organoid chamber (hvCOC) mini-hearts. (A) Representative images of time-matched (Day 10) hvCOC under control conditions and with combined TGF-β1/ET-1 treatment. (B) Change in diastolic pressure versus change in diastolic area for control (black) and TGF-β1/ET-1-treated (red) hvCOC subject to 0-200 µL hydrostatic loading. Slope of the graph reflects diastolic stiffness. (C) Dot plot of hvCOC diastolic stiffness at 100-µL hydrostatic loading for control (black circles) and TGF-β1/ET-1-treated hvCOC (red circles). (D) Representative pressure-area loops for control (black) and TGF-β1/ET-1-treated (red) hvCOCs at 2.0-Hz pacing. (E) Corresponding changes in pressure (left) and area (right) over time for control (black) and TGF-β1/ET-1-treated (red) hvCOCs. (F) Dot plots for developed pressure, stroke area, stroke work and ejection fraction for control (black) and TGF-β1/ET-1-treated (red) hvCOCs paced at 2 Hz field stimulation. (B), (C), and (F) n=8-13 mini-hearts; mean ± SEM; Student’s two-tailed t-test. *p<0.05.

### HFpEF induction applied to hvCOC mini-heart exhibits elevated diastolic chamber stiffness with preserved ejection fraction

The combined TGF-β1/ET-1 administration was then assessed for the ability to confer HFpEF phenotypic characteristics to the hvCOC mini-heart model, created from the same healthy hPSC-derived hvCMs and fibroblasts as previously detailed^22, 26^ to obtain a three-dimensional (3-D) electro-mechanically coupled, fluid-ejecting ventricular chamber, in a bioreactor system enabling automated multi-heart measurements. **Figure 2A** shows representative images of time-matched control and TGF-β1/ET-1-treated hvCOCs, in which the diastolic area was not significantly different in HFpEF versus Control (p = 0.244, not shown). **Figures 2B** and **C** show a leftward-shifted diastolic pressure-area relationship, and significantly higher passive chamber stiffness in TGF-β1/ET-1-treated versus control hvCOCs, respectively (n=8-13, *p<0.05). Despite the increased stiffness, pressure-area tracings were similar (**Fig. 2D, E**), with no significant changes in developed pressure, stroke area, stroke work or ejection fraction observed between Control and TGF-β1/ET-1-treated hvCOCs (**Fig. 2 E-F**). Therefore, TGF-β1/ET-1-treated hvCOC displayed phenotypes consistent with HFpEF characteristics. Similar results were obtained using hvCOC mini-hearts created from a second independent hPSC cell line, supporting the robustness of our HFpEF model (**Supplemental Fig. 3**).

### Transcriptomic signatures of HFpEF patients and human mini-Heart models identify sarcoplasmic reticulum (SR) Ca^2+^-ATPase pump (SERCA2a) as a human therapeutic target

Next, we performed transcriptomic and bioinformatic analyses of differentially expressed genes (DEGs) from bulk RNA sequencing of induced-HFpEF hvCTS and hvCOC mini-heart models versus respective controls, and compared these to published DEGs from right ventricular septal biopsies from HFpEF patients versus organ donor controls^8^ (**Fig. 3**). Venn diagrams showed that 29% (501/1748) of DEGs from the HFpEF-hvCTS model, and 33% (446/1351) of DEGs from the HFpEF-hvCOC model, overlapped with DEGs from HFpEF patients (**Fig. 3A**). Ingenuity Pathway Analysis further revealed that the HFpEF-hvCOC mini-ventricle model was more similar to human HFpEF patients than the HFpEF-hvCTS tissue strip model based on hierarchical clustering of enriched canonical pathways, disease and biofunctions, and cardiotoxicity functions (**Fig. 3B**). Interestingly, the Calcium Signaling pathway was differentially activated in both human HFpEF patients and in the HFpEF-hvCOC model, but not in the HFpEF-hvCTS model, relative to their respective controls (Fisher’s exact test P-values: human HFpEF; p=0.008, hvCOC; p=0.01 and hvCTS; p=0.10) (**Fig. 3B**). Hierarchical clustering of DEGs in the Calcium Signaling Pathway (**Fig. 3C**) showed that the ATP2A2 gene (encoding for SERCA2a) was downregulated in HFpEF patients as well as engineered HFpEF mini-heart models compared to their respective controls (**Fig. 3D**). Of note, ATP2A2 was one of the most highly expressed genes in the gene ontology (GO) term “Regulation of ATPase-coupled calcium transmembrane transport activity” in normal donor hearts as well as control engineered hvCTS and hvCOC models (**Fig. 3E**). These transcriptomic and bioinformatic results were consistent with the phenotypes observed in HFpEF patients, thereby identifying restoration of SERCA2a as a target candidate for mitigating the HFpEF disease traits.

**Figure 3.**
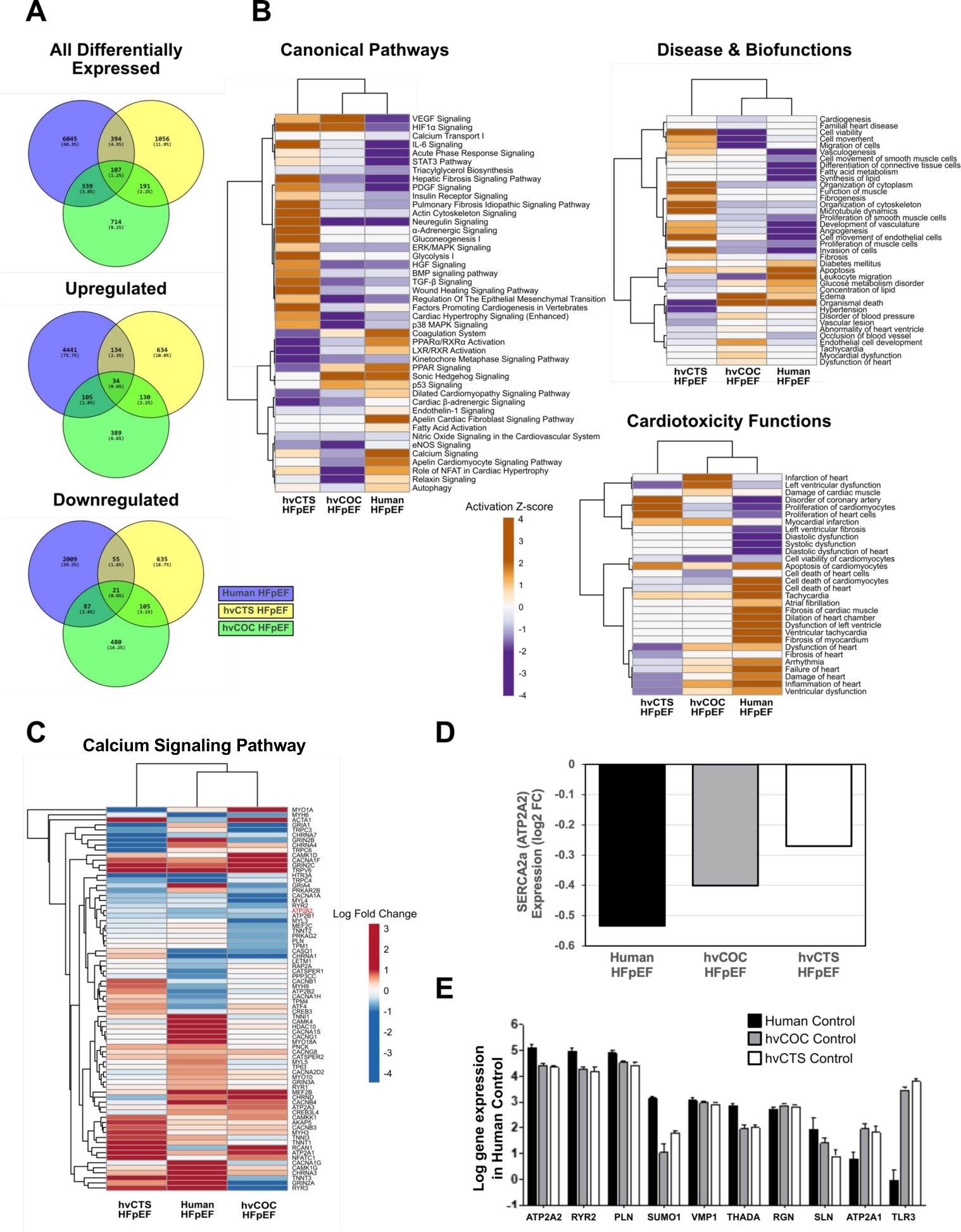
Transcriptomic and bioinformatic analyses of HFpEF patient heart tissue (n = 41) compared to hvCTS (n = 7) and hvCOC (n = 5) mini-Heart models, revealing that SERCA2a was downregulated in HFpEF. (A) Venn diagrams of all differentially expressed genes (DEGs), including upregulated and downregulated DEGs, from the HFpEF-hvCTS and HFpEF-hvCOC heart models versus respective controls, as well as DEGs from HFpEF patients versus donor hearts. (B) Hierarchical clustering of a selection of enriched canonical pathways, disease and biofunctions and cardiotoxicity functions based on Ingenuity Pathway Analysis (IPA) of DEGs from the HFpEF-hvCTS and HFpEF-hvCOC mini-heart models versus respective controls, as well as from HFpEF patients versus donor hearts (Fisher’s exact test P-value ≤ 0.05 in at least one contrast). Heatmaps are colored by IPA activation Z-scores (orange: predicted activation, purple: predicted inhibition). (C) Hierarchical clustering of genes in IPA Calcium Signaling Pathway. Heatmaps are colored by log2(FC) values from differential expression analysis of the HFpEF-hvCTS heart model, HFpEF-hvCOC heart model, and HFpEF patients versus respective controls. (D) The SERCA2a (ATP2A2) DEG was downregulated in HFpEF patients and in engineered HFpEF-hvCTS and HFpEF-hvCOC mini-heart models compared to respective controls. (E) Bar graph comparing the ten highest expressed genes in the gene ontology (GO) term “Regulation of ATPase-coupled calcium transmembrane transport activity” in donor hearts and control engineered hvCTS and hvCOC models (mean±SD, n=3 tissues).

### Dosage optimization – AAV transduction of human mini-heart models is highly titre- and time-dependent

As a next step, we investigated the titer- and time-dependencies of recombinant adeno-associated vector type 1 (AAV1) transduction of hvCMs. Green Fluorescent Protein (GFP) expression increased with increasing AAV1-GFP dosage at Day 14 post-transduction (**Fig. 4A**). The percentage of GFP-positive hvCMs also gradually increased over time after AAV1-GFP transduction (**Fig. 4B**), reaching a plateau that was dependent on the viral titer (i.e., viral genomes per cell, vg/cell). Accordingly, **Figure 4C** shows that SERCA2a gene expression in hvCMs after 14 days of transduction with AAV1-SERCA2a was similarly titer- and time-dependent (one-way ANOVA; *p<0.05), suggesting that optimal titers will be critical for the success of cardiac gene therapies in human patients.

**Figure 4.**
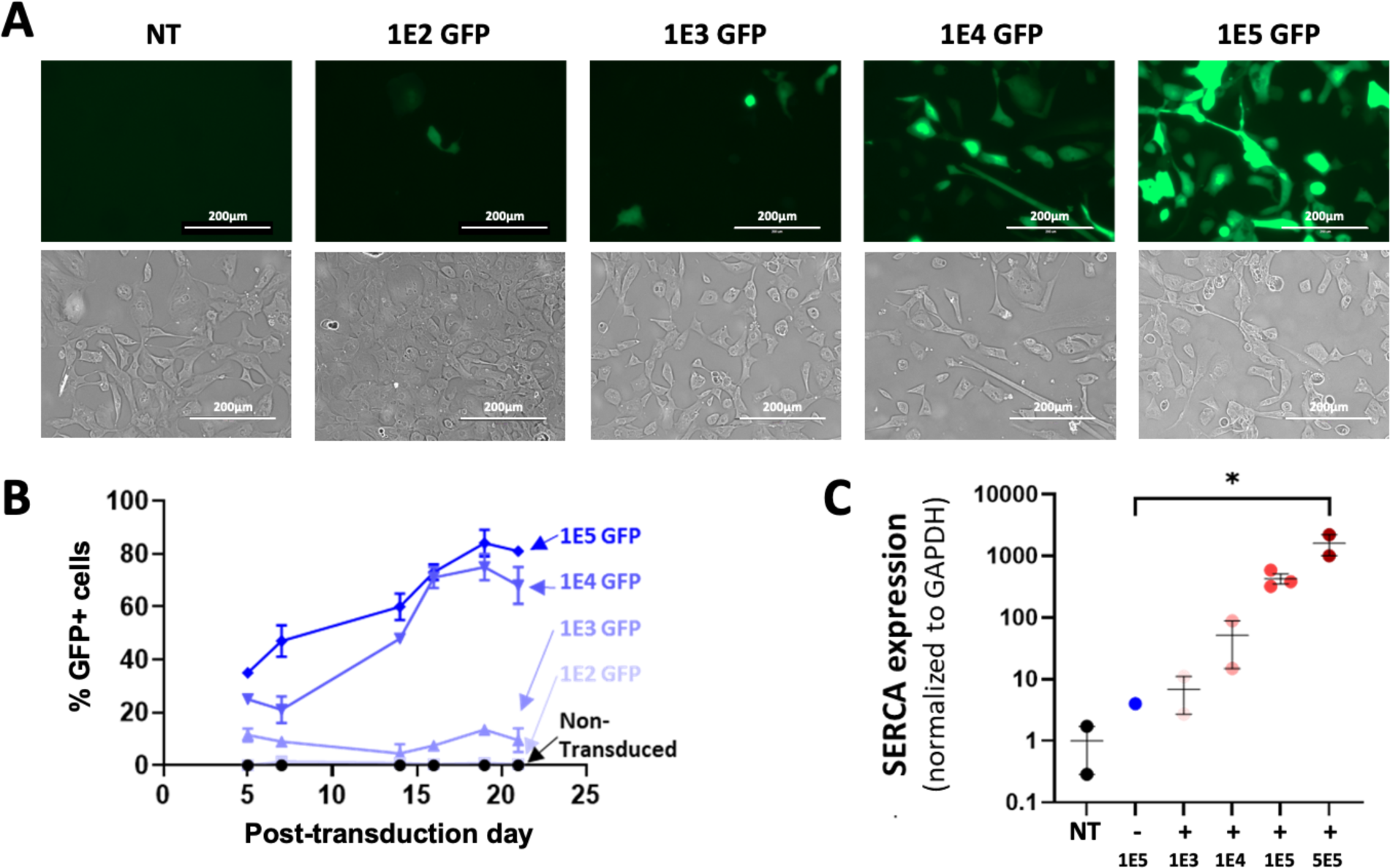
Titer- and time-dependence of AAV transduction. (A) Representative fluorescence and brightfield images of hvCM monolayers after 14 days of AAV1-GFP transduction with different viral genome (vg) concentrations, ranging from 1×10^2^ vg/cell (1E2 GFP) to 1×10^5^ vg/cell (1E5 GFP). (B) The effect of virus titer on the percent of GFP-expressing cells versus time after AAV1-GFP transduction. (C) SERCA gene expression level in hvCMs, by RT-qPCR, after 14 days of AAV1-GFP (-) or AAV1-SERCA2a (+) transduction, with titers as shown. NT is non-transduced control. GAPDH is housekeeping gene. n =1-3 plates; mean ± SEM; one-way ANOVA; *p<0.05.

### AAV1-SERCA2a rescues the disease phenotype in HFpEF hvCTS tissue strips and hvCOC mini-hearts

Based on the above data, we selected a titer of 1×10^5^ vg/cell and 14 days post-transduction for studying the effects of AAV1-SERCA2a-mediated overexpression in HFpEF-hvCTS. **Figure 5A** shows representative images of time-matched HFpEF-hvCTS after AAV1-GFP (control) and AAV1-SERCA2a treatments. **Figure 5B** shows that in representative HFpEF hvCTS tracings, AAV1-SERCA2a transduction hastened the slowed contraction and relaxation kinetics compared to the AAV1-GFP control. This is corroborated in **Figure 5C**, where AAV1-SERCA2a treatment improved the abnormal kinetics of the HFpEF hvCTS by significantly shortening the diseased contraction and relaxation times toward non-transduced (NT) Control tissue values (n=5-8, p<0.05), while active and passive force were increased. On average, the data for AAV1-GFP and NT HFpEF hvCTS were very similar.

**Figure 5.**
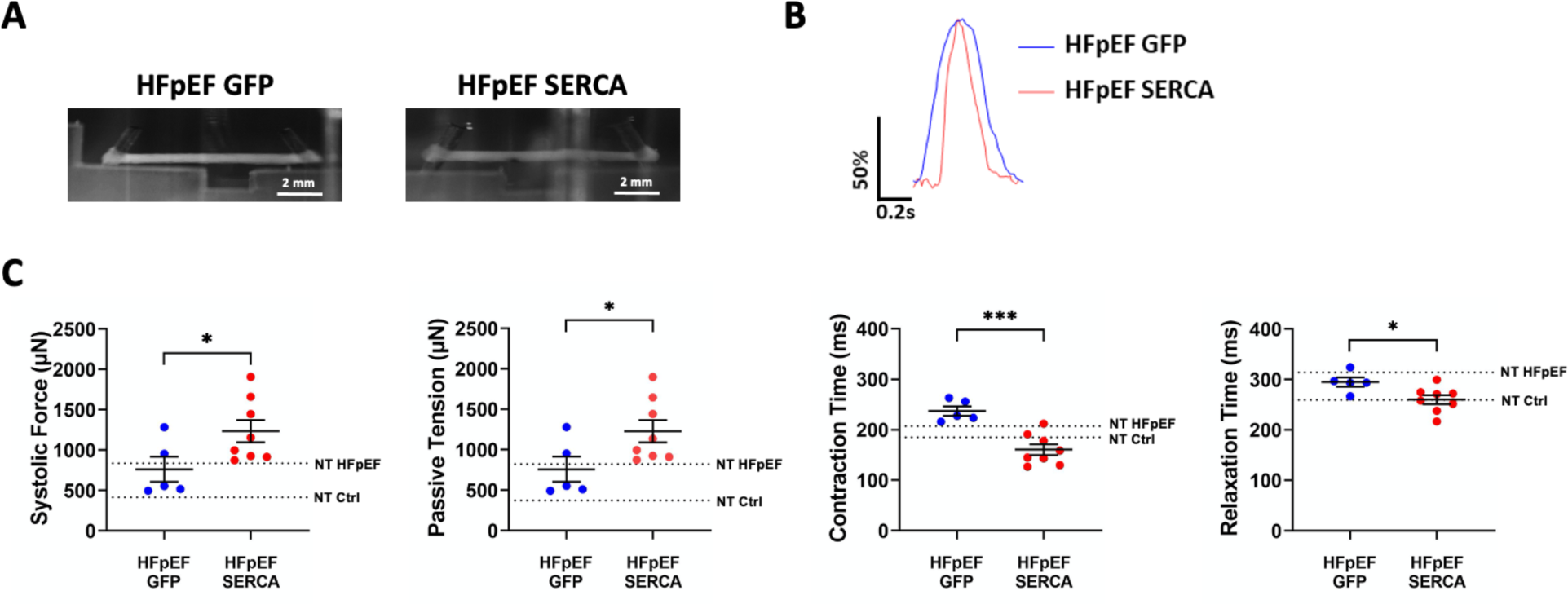
AAV1-SERCA2a improves HFpEF phenotype in hvCTS. (A) Representative images on Day 10 of HFpEF-hvCTS transduced with AAV1-GFP or AAV1-SERCA2a. (B) Representative normalized single-twitch force tracings of HFpEF-hvCTS with AAV1-GFP (blue line) or AAV1-SERCA2a (red line) at 1.5-Hz pacing. (C) Dot plots for the systolic force, passive tensile force, contraction time and relaxation time as recorded by post-tracking measurements during pacing at 1.5 Hz field stimulation. Data are shown as mean ± SEM for HFpEF-hvCTS with AAV1-GFP (blue circles) and HFpEF-hvCTS with AAV1-SERCA2a (red circles). Dashed lines indicate mean values for non-transduced (NT) healthy controls and NT HFpEF hvCTS samples created from the same batch of cells. n =5-8 tissues, *p<0.05 and ***p<0.001 by Student’s two-tailed t-test.

We further examined the effects of AAV1-SERCA2a treatment in the hvCOC mini-heart model of HFpEF (**Fig. 6**). For a balance between successful chamber compaction and efficient transduction, 1×10^4^ vg/cell was used. As anticipated from our HFpEF hvCTS experiments, the related measures of pump function including developed pressure, stroke area, stroke work, and ejection fraction of AAV1-GFP- and AAV1-SERCA2a-treated HFpEF hvCOC mini-hearts were not significantly affected (**Fig. 6D-E**). By contrast, AAV1-SERCA2a treatment significantly shortened area and pressure contraction times (**Fig. 6E**). Taken together, these findings provide evidence of hastened contraction kinetics in the HFpEF-hvCOC mini-heart model treated with AAV1-SERCA2a, though the phenotype correction is clearer in the hvCTS model.

**Figure 6.**
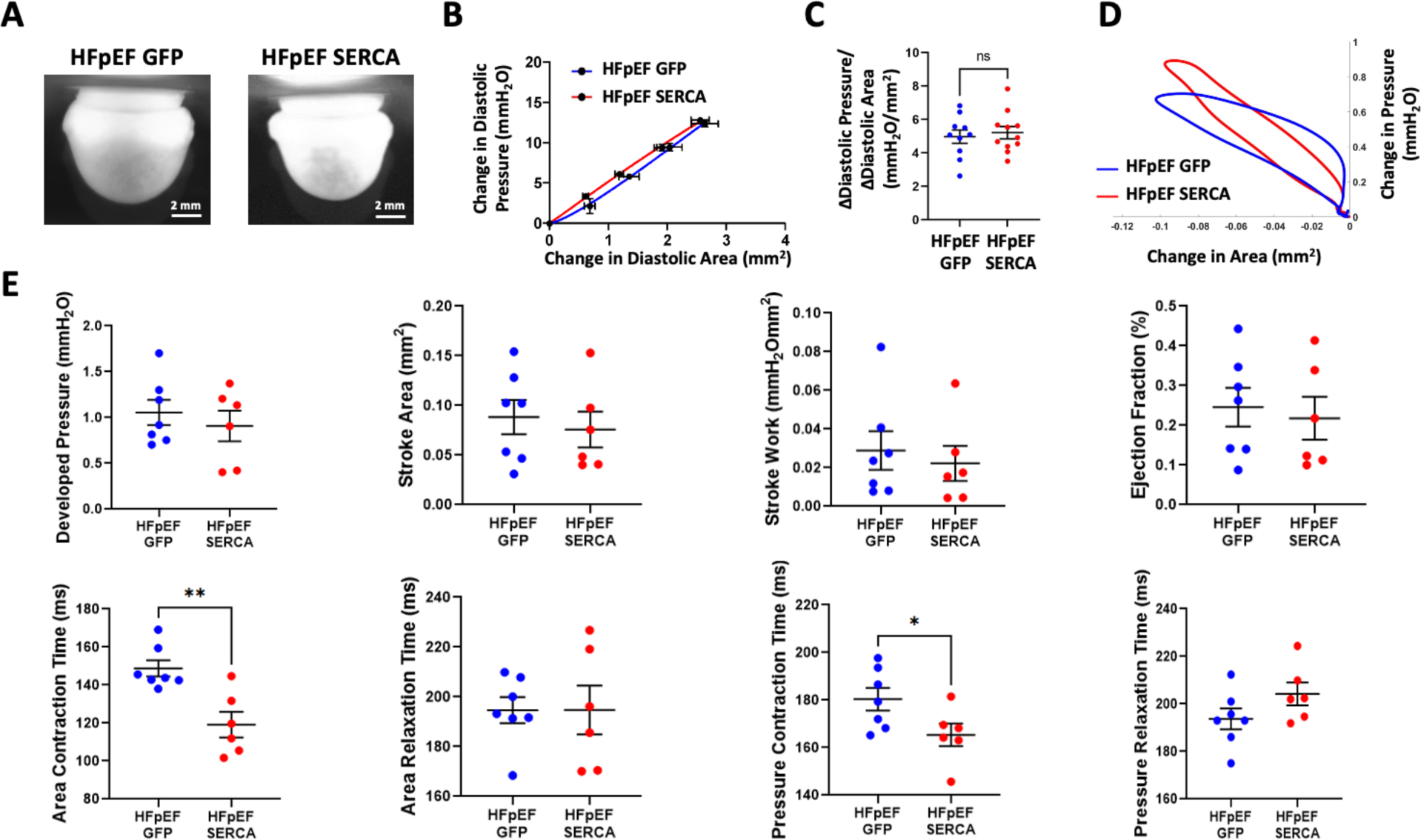
AAV1-SERCA2a improves HFpEF phenotype in hvCOC. (A) Representative images of HFpEF-hvCOC transduced with AAV1-GFP (top) or AAV1-SERCA2a (bottom), imaged on Day 31 post-fabrication. (B) Diastolic pressure vs. area plot for AAV1-GFP-treated (blue) and AAV1-SERCA2a-treated (red) HFpEF hvCOC models subject to 0-200µL hydrostatic loading. (C) Dot plot of HFpEF hvCOC diastolic stiffness at 100-µL hydrostatic loading for AAV1-GFP control (blue) and AAV1-SERCA2a treatment (red). (D) Representative pressure-area loops of HFpEF hvCOC at 2.0-Hz pacing, treated with AAV1-GFP (blue) or AAV1-SERCA2a (red). (E) Dot plots of developed pressure, stroke area, stroke work, and ejection fraction (top row) as well as area and pressure contraction and relaxation times (bottom row), for HFpEF-hvCOC treated with AAV1-GFP (blue circles) or with AAV1-SERCA2a (red circles). (B-C) n=10-11 mini-hearts, mean ± SEM; Student’s two-tailed t-test. (E) n=6-7 mini-hearts, mean ± SEM; *p<0.05, **p<0.01 by Student’s two-tailed t-test.

## Discussion

Three-dimensional bioengineered tissue constructs have been developed using cardiomyocytes derived from healthy hPSCs. Despite the sub-physiologic levels of function typical of engineered tissue constructs for *in vitro* studies, the hvCTS and hvCOC human mini-heart models used in this study have proven effective for modeling different cardiovascular disease states and testing of drugs and biologics.^17, 18, 22, 23^ The value of such model systems has been reinforced by the recent FDA Modernization Act 2.0, which promotes the use of alternatives to animal testing for more predictive pre-clinical studies.^21^ Toward this goal, the *in vitro* human HFpEF heart models described herein have been constructed using multiple hPSC cell lines, and rigorously validated using a range of phenotypic assessments recapitulating key characteristics of the disease, including favorable comparison with HFpEF patient transcriptomic data. Challenges persist for *in vitro* model systems to reproduce complex comorbidities such as diabetes and obesity that often accompany patients with HFpEF,^7^ yet salient features of the cardiac phenotype may be adequately represented in engineered mini-heart models. Considering the diverse nature of HFpEF, the new hvCTS and hvCOC models here may represent a more specific patient subpopulation.

In contrast to prior cardiac disease modeling approaches focused on genetic mutations,^20, 42, 43^ this study demonstrates the use of biologics for induction of disease without requiring patient-specific human induced pluripotent stem cells. The results supported SERCA2a as a human-relevant therapeutic target for HFpEF, and further demonstrated the improvement of characteristic HFpEF phenotypes by AAV1-SERCA2a treatment in human mini-hearts, which contributed to an ongoing first-in-human gene therapy clinical trial for HFpEF (Modulation of SERCA2a of Intra-myocytic Calcium Trafficking in Heart Failure with Preserved Ejection Fraction (MUSIC-HFpEF, NCT06061549)). These models may be suitable for assessing a variety of candidate agents acting on the myocardium to treat the HFpEF condition, including viral vectors and small molecules, and we anticipate that such models will play an increasingly important role for future regulatory decisions.

In mammalian hearts, HFpEF is associated with impaired cardiac relaxation, or lusitropy, which is evident both at rest and during exercise in the majority of HFpEF patients.^6, 44^ Cardiac myocytes isolated from HFpEF patients and experimental models exhibit prolonged relaxation, diminished contraction velocity, a decrease in β-adrenergic response, and increased myocardial stiffness.^45, 46^ The abnormalities in cardiac relaxation in experimental models of HFpEF have been attributed to deficiency in the content and activity of SERCA2a,^12, 47-49^ which decreases the rate of Ca^2+^ uptake into the SR during relaxation,^12, 13, 47, 48^ resulting in prolonged contraction. Furthermore, SERCA2a protein levels were found to be significantly decreased in senescent human myocardium that is characterized by HFpEF,^50^ which was associated with myocardial function impairment at baseline that was exacerbated by hypoxic conditions and higher heart rates.^51-53^ When heart rate increases, calcium that is slow to dissociate from troponin-actin-myosin complexes after contraction creates a “rigor-like” state that may contribute to increased diastolic stiffness. In addition, the slow re-uptake correlates directly with the relaxation phase of pressure. In experimental models of HFpEF (e.g., Dahl salt sensitive rats, spontaneously hypertensive, aortic banding, diabetic, aged), SERCA2a expression has been found to vary from no change to a decrease even though intracellular resting calcium has been found to be invariably elevated in these HFpEF models.^12, 47-49, 54-57^ Thus, the consequence of slowed calcium re-uptake has implications not only for myocardial relaxation but also contributes to the development of diastolic stiffness, key features in the pathophysiology of HFpEF.

An approach to restore intracellular calcium homeostasis is to enhance calcium uptake by SERCA. This can be achieved by modulating SERCA activity or increasing expression of SERCA pumps. We have previously shown that in HFpEF animal models characterized by abnormal lusitropy and a decrease in SERCA2a, delivery of SERCA2a by gene transfer restored relaxation parameters such as -dP/dt, the left ventricular time constant of isovolumic relaxation, tau, and the passive stiffness of the left ventricle to normal levels.^12, 13, 47-49^ These results establish that viral delivery of SERCA2a in HFpEF hearts can improve relaxation parameters.^13^ In other studies, the Otsuka-Long-Evans Tokushima Fatty rat model of spontaneous non-insulin-dependent type II diabetes mellitus, which is characterized by diastolic dysfunction and is associated with decreased SERCA2a expression, was used as a model of HFpEF. In multiple studies using short-term expression of SERCA2a, oxygen consumption and relaxation parameters were restored to normal levels in these HFpEF models.^12, 47, 48^ Our current work provides the first direct link to human relevance.

The experiments reported herein establish the hvCTS and hvCOC mini-heart models of HFpEF in human species-specific diseased heart tissues. The data further suggest treatment methods for HFpEF involving increasing, or restoring, intracellular calcium homeostasis in cardiomyocytes by increasing or enhancing calcium uptake by SERCA to ameliorate or reverse the slowed contraction and relaxation kinetics in HFpEF, which is translatable to humans. Indeed, our new MUSIC-HFpEF trial using AAV1-SERCA2a gene therapy is the only current human clinical trial targeting relaxation in HFpEF, and inclusion of data from this study using the same GMP-grade viral vector contributed to our successful Investigational New Drug (IND) and Fast Track Designation (FTD) applications to the U.S. FDA.

The CUPID study was a first-in-human gene therapy trial for HF that also focused on improving SERCA function. In Phase 1 (CUPID-1), HFrEF patients randomized to the highest dose of a single intra-coronary infusion of AAV1-SERCA2a showed less deterioration in 6-minute walk distance, peak VO_2_ and NT-proBNP levels after 6 months.^58-60^ A larger Phase 2 study (CUPID-2), testing a higher dose (10^13^ viral genome particles of AAV1-SERCA2a per patient), confirmed the safety of this approach,^61, 62^ but showed no effect on the primary outcome of HF hospitalization or decompensated HF. Follow-up analysis of available patient cardiac tissue revealed transduction efficiency far below doses that were effective in preclinical models, which likely explained the neutral outcome.^63^ Also, the CUPID-2 trial’s strategy was to increase SERCA2a activity as a means of improving contractile dysfunction characteristic of HFrEF; the effect on cardiac relaxation was never assessed.

In the ongoing MUSIC-HFpEF trial, increasing SERCA2a activity or expression is expected to be an effective therapeutic target in HFpEF whereby improved lusitropy would likely allow tolerance of higher heart rates and increase diastolic filling of the ventricles, improving exercise capacity and peak VO_2_, and a decrease in left ventricular filling pressures. Despite experimental evidence that SERCA2a gene transfer improves cardiac function in HFrEF, and functional deficiencies of SERCA2 have been well validated in the context of HFrEF, there are mixed reports regarding the protein levels of SERCA2a in humans.^64-76^ However, the current study data clearly indicate that SERCA2a represents a relevant target for human HFpEF. Indeed, the human mini-heart models of HFpEF described herein demonstrate improvements in myocardial kinetics with AAV1-SERCA2a treatment.

Our human mini-heart models also provide a relevant platform for dosage optimization, considering that results from animal experiments regarding effectiveness of cardiac gene transfer do not predict human myocardial transduction efficiency.^77^ Without positive transduction, gene therapy cannot be effective; the above data strongly indicated that optimal titers are critical for the success of cardiac gene therapies in humans, in addition to other considerations such as delivery method, vector serotype and tropism, and potential side effects.^78^ Considering there are 2 to 3 billion cardiomyocytes in the adult human heart, the optimized viral titer of 1×10^4^ vg/cell used in the hvCOC mini-heart model of HFpEF translates to a dosage of 2-3×10^13^ vg per patient heart; this informed the approved dosage of 3 x10^13^ vg/patient to be delivered via intracoronary injection of HFpEF subjects in the MUSIC-HFpEF trial. Together with the observation reported herein that higher viral titers were beneficial in transducing hvCMs, it is expected that higher doses of AAV1-SERCA2a than were used in CUPID-2 will provide an effective treatment for patients with HFpEF, based on precise characterization of myocardial relaxation and filling pressures during exercise before and after treatment. Encouraging early results from the MUSIC-HFpEF trial are indicating improvements in NYHA HF classification, lowering of pulmonary capillary wedge pressure during exercise, and reduced levels of circulating NT-proBNP, supporting efficacy of the AAV1-SERCA2a treatment.

In conclusion, the reported HFpEF-hvCTS and HFpEF-hvCOC mini-heart models are uniquely of human origin in the disease setting (HFpEF), and amenable to sophisticated phenotypic measurements, as in the above experiments, demonstrating the ability to recapitulate characteristic phenotypes seen in HFpEF patients. Using the SERCA2a transgene as an example, we also demonstrated how the human mini-Heart HFpEF models are used to identify novel druggable targets followed by therapeutic screening for cardiac functional improvements. The versatile HFpEF models can be further custom-tailored, as needed, for discovering additional novel targets and screening therapeutics; the human mini-heart model is also helpful for titrating dosages in human trials. Therefore, our approach is aligned with guidelines of the recent FDA Modernization Act 2.0 in support of alternatives to animal testing. Our study also underscores the value of looking for common patterns that may arise from diverse yet complementary experimental disease models that each have strengths and limitations. Overall, the preclinical human HFpEF models described herein are shown to be useful in investigating disease mechanisms, and for facilitating the identification of novel druggable targets and screening of therapeutics by enabling species-specific predictions of the clinical effectiveness and safety of new medicines in advance of human trials.

## Author Contributions

KDC, RJH, RAL wrote the manuscript and supervised the study. AOTW, SYM, CC, AW performed experiments, collected and analysed data, and prepared figures. EGR, WK, JC, AC, DKL assisted with data analysis and interpretation. AOTW, DKL assisted with editing the manuscript. KJ, QDW supervised aspects of the study, assisted with data interpretation, and reviewed and edited the manuscript.

## Acknowledgements

The authors acknowledge technical support from Daniel Jachimowicz, Julia Lindgren, Karl Nordström with RNA sequencing and data pre-processing. We also thank Jeff Rudy from Sardocor, Medera Inc., for generously providing the GMP-grade AAV1-SERCA2a drug product used in this study.

## Data Availability

Data supporting the findings of this study are available within the paper and its Supplementary Information. RNA sequencing data will be made available online prior to publication of this study. Raw data will be made available upon request to the corresponding authors.

## Code Availability

In-house custom code was developed and implemented using LabVIEW (v. 2018, National Instruments) for the collection and MATLAB (v. 2020-2022, MathWorks) for the analysis of the tissue data presented in this manuscript. Interested researchers can request access to the code by contacting the corresponding authors. Access to the code may be subject to certain restrictions or licensing agreements.

