## Supplementary Figures for "Reversal of contractile defects by mediating calcium homeostasis in human mini-heart models of heart failure with preserved ejection fraction (HFpEF) leads to first-in-human gene therapy clinical trial"

2., Research and Early Development, Cardiovascular, Renal and Metabolism (CVRM), BioPharmaceuticals R&D, AstraZeneca, Gothenburg, Sweden

3. Sardocor, Medera Inc, Boston, MA, USA

*co-corresponding authors

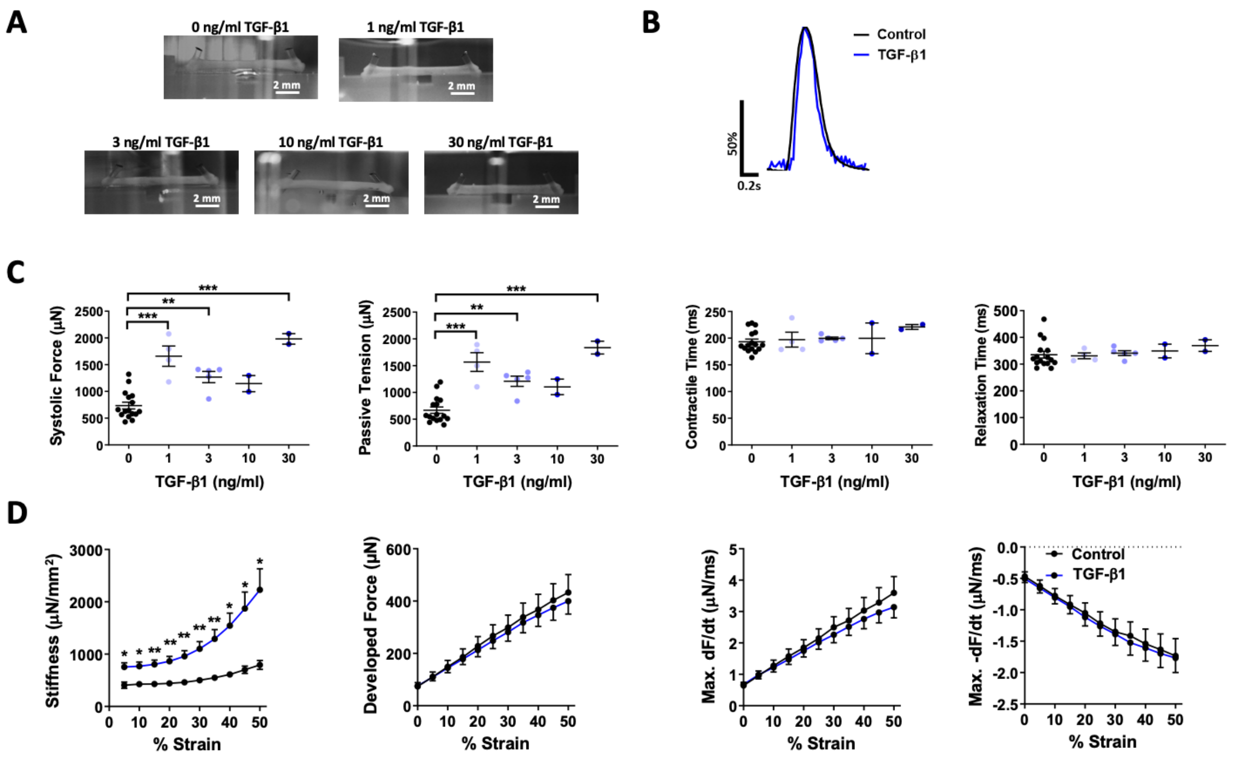


**Supplemental Figure 1.** The effects of TGF-β1 on hvCTS. (A) Representative images of hvCTS treated with 0–30 ng/ml TGF-β1, in the passive diastolic state. (B) Representative post-tracking normalized force tracing of hvCTS without TGF-β1 (Control; black line) and 1 ng/ml TGF-β1 (blue line) at 1.0-Hz pacing. (C) Systolic force, passive tensile force, contraction time and relaxation time measured by post-tracking. 0 ng/ml TGF-β1 (black circles); 1 ng/ml TGF-β1 (light blue circles); 3 ng/ml TGF-β1; 10 ng/ml TGF-β1; or 30 ng/ml TGF-β1 (dark blue circles). n =2-16, mean ± SEM; One-Way ANOVA followed by Dunnett's multiple comparison test. (D) hvCTS stiffness, developed force, max +dF/dt and max -dF/dt versus percent strain for the Control (black) and 1 ng/ml TGF-β1 (blue) groups measured using the isometric muscle bath system. n=12-16; mean ± SEM; Two-Way ANOVA followed by Sidak's multiple comparison test. *$p<0.05$, **$p<0.01$ and ***$p<0.001$.

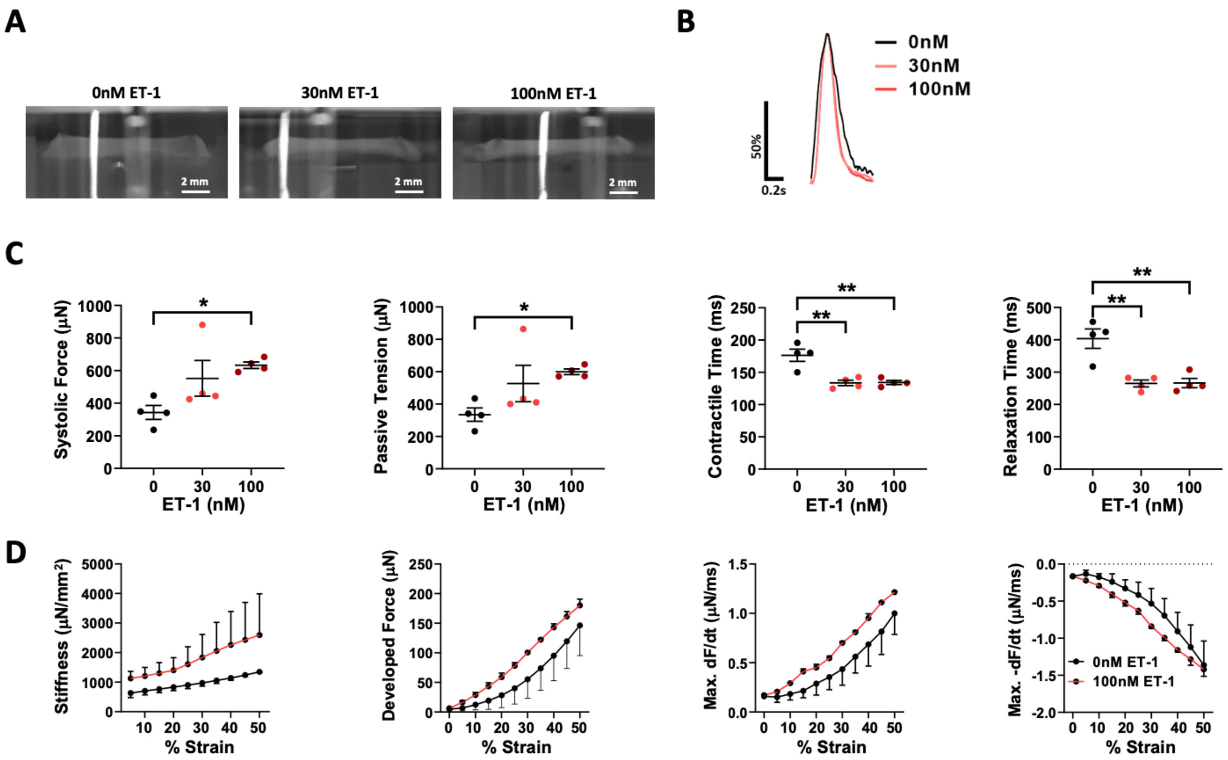


**Supplemental Figure 2.** The effects of ET-1 on hvCTS. (A) Representative images of hvCTS treated with 0–100 nM ET-1, in the passive diastolic state. (B) Representative post-tracking normalized force tracing of hvCTS. 0 nM ET-1 (black line); 30 nM ET-1 (light red line); 100 nM ET-1 (dark red line) at 1.0-Hz pacing. (C) Systolic force, passive tensile force, contraction time and relaxation time measured by post-tracking. 0 nM ET-1 (black circles); 30 nM ET-1 (light red circles); 100 nM ET-1 (dark red circles). n = 4 tissues, mean ± SEM; One-Way ANOVA followed by Dunnett's multiple comparison test. (D) hvCTS stiffness, developed force, max +dF/dt and max -dF/dt versus percent strain for the control (black) and 100 nM ET-1 (red) groups measured using the isometric muscle bath system. n = 2 tissues; mean ± SEM; Two-Way ANOVA followed by Sidak's multiple comparison test. *$p<0.05$ and **$p<0.01$.

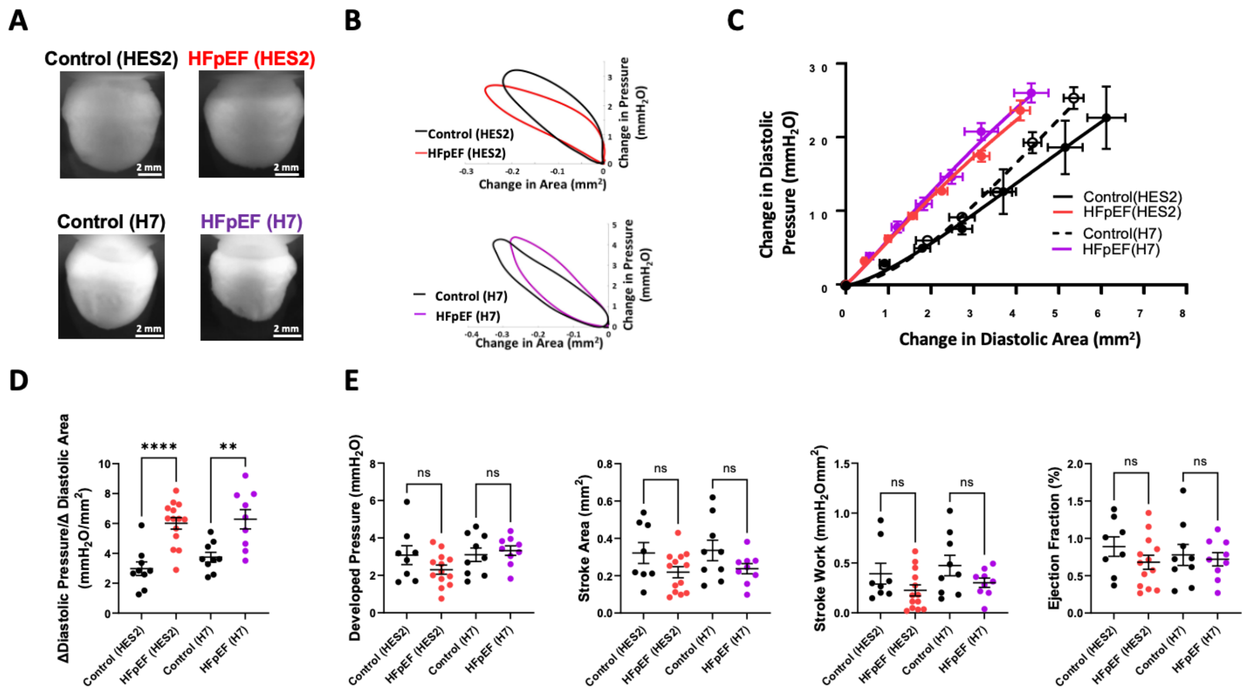


**Supplemental Figure 3.** Consistency of hvCOC HFpEF mini-heart models using multiple hPSC cell lines. (A) Representative images of time-matched (Day 10) hvCOC for Control and HFpEF induced with combined TGF-β1/ET-1 treatment, using both the HES2 (top) and H7 (bottom) hPSC cell lines. (B) Representative pressure-area loops at 1.5-Hz pacing for Control (black) and HFpEF (colored) hvCOCs using HES2 (top) and H7 (bottom) hPSC cell lines. (C) Change in diastolic pressure versus change in diastolic area for Control (black) and HFpEF (colored) hvCOC subject to 0-200 μL hydrostatic loading, using HES2 and H7 hPSC lines as indicated in the key; slope of the graph reflects diastolic stiffness. (D, E) Dot plots for hvCOC diastolic stiffness at 100-μL hydrostatic loading, developed pressure, stroke area, stroke work and ejection fraction for Control (black) and HFpEF (colored) hvCOCs paced at 2 Hz, using HES2 and H7 hPSC cell lines as indicated. (C)-(E) n=8-13 mini-hearts, mean ± SEM; Student's twotailed t-test. **$p<0.01$ and ****$p<0.0001$. Note that some results for HES2 were also shown in main Figure 2.
